# Novel regulators of zebrafish glomerular development

**DOI:** 10.64898/2026.10.27.762292

**Authors:** Omodasola Adekeye, Blanca de Juan Mora, Ritu Tomar, Olaleye Olajuyin, Clemence Rouzier, Nicole Rainford, Brian Hartman, Daemon Dikeman, Celeste Nobrega, Joel Graber, Caramai Kamei, Iain A. Drummond

## Abstract

Podocyte cell differentiation in coordination with specialized endothelial cells forms the glomerulus, the kidney blood filter. Genetic analysis of animal models and human affected individuals has identified over 50 susceptibility genes that, when mutated, cause filtration defects or proteinuria. To explore the potential for new genetic pathways or components in glomerular development, we generated a transcriptome of micro-dissected three-day-post-fertilization zebrafish larval pronephric glomeruli and conducted a CRISPR-Cas9 G0 mutagenesis screen of less well characterized genes that showed glomerulus-enriched expression. Differential gene expression comparing glomeruli to whole larvae revealed a significant overlap of zebrafish glomerular-enriched gene expression with the human podocyte transcriptome. CRISPR-Cas9 G0 mutagenesis of twelve genes identified eight potential new causes of proteinuria in larval zebrafish based on glomerular leakage of rhodamine-labelled, 500 kDa high molecular weight dextran. Novel causes of proteinuria include regulators of small GTPases (*fam49ba/cyrib, rasgrp3*), cell adhesion/phosphatidylinositol metabolism (*ptprq*), and an upstream regulator of the hippo pathway (*frmd6*). Our glomerular transcriptome and screening results open new opportunities to explore glomerular cell differentiation and function.

## Introduction

The glomerulus is the filtration unit of the kidney, responsible for maintaining physiological homeostasis and ensuring the retention of essential plasma proteins during filtration (Arif and Nihalani, 2013; Brenner et al., 1978; Itoh et al., 2014). Its highly specialized structure consists of fenestrated endothelial cells, a basement membrane, and podocytes, an epithelial cell with elaborate foot processes that interdigitate and are bridged by membrane-spanning proteins called the slit diaphragm (Pavenstädt et al., 2003; Perico et al., 2016). Disruption of glomerular function is marked by the loss of protein in the urine, known as proteinuria, which is a clinical hallmark of glomerular disease and a strong predictor of progression to end-stage renal disease (Daehn and Duffield, 2021; Haraldsson et al., 2008; Wiggins, 2007)

To date, over 50 genes have been linked to glomerular maldevelopment and disease, including those encoding podocyte slit diaphragm proteins (e.g., NPHS1, NPHS2), components of the actin cytoskeleton (e.g., ACTN4), signaling molecules (e.g., TRPC6, PLCε1), and transcription factors crucial for podocyte differentiation (e.g., WT1) (Kopp et al., 2020; Preston et al., 2019). Given the complexity of podocyte cell structure and the requirement for integrated development with the vasculature, it is likely that additional genes and pathways involved in glomerular development remain to be discovered.

The zebrafish larval pronephric glomerulus is a model to study the function of glomerular genes using forward and reverse genetic approaches, chemical screens, and high-resolution imaging (Hanke et al., 2013; Kramer-Zucker et al., 2005; Lee et al., 2022). The zebrafish embryonic kidney consists of a pronephric tubule on each side and a single glomerulus with a glomerular filtration barrier (Drummond et al., 1998), which is similar to that in mammals, including humans. Pronephric glomerular morphogenesis begins at 24 hpf with podocyte cell differentiation and by 96 hpf, a selective filtration barrier is formed (Drummond, 2005; Drummond et al., 1998; Kramer-Zucker et al., 2005; Nishibori et al., 2011). Beyond the structural resemblance of the zebrafish embryonic kidney to higher vertebrates, many of its gene expression features are similarly conserved (Drummond and Davidson, 2010; Ebarasi et al., 2011; Kroeger and Wingert, 2014), indicating that investigating the function of glomerulus-enriched genes in zebrafish offers a strategy for identifying novel pathways that contribute to glomerular development.

In this study, we combined glomerulus-specific bulk RNA seq of larval zebrafish glomeruli with a CRISPR-Cas9-based mutagenesis screen to identify new regulators of zebrafish glomerular development. Functional analysis of mosaic mutants revealed novel causes of proteinuria, including genes involved in small GTPase regulation and phosphatidylinositol metabolism. We report previously unrecognized regulators of glomerular morphogenesis and provide a foundation for future mechanistic studies of podocyte cell differentiation.

## Materials and methods

### Zebrafish husbandry

Adult zebrafish were maintained at 28.5°C in tanks with constant water flow and fed twice a day, according to standard protocols (Westerfield, 2000). Larvae were raised at 28°C in E3 medium (≤ 100 per dish), with medium changes at 2 dpf and daily removal of dead larvae. For procedures, larvae were anesthetized in 0.02% tricaine methanesulfonate (MS 222; Sigma-Aldrich) and euthanized in 0.2% tricaine methanesulfonate. All studies adhere to the National Institutes of Health Guide for the Care and Use of Laboratory Animals and were approved by the Mount Desert Island Biological Laboratory Institutional Animal Care and Use Committee.

### Glomerulus isolation and sequencing

A cross of transgenic *Tg(podocin:gal4) X Tg(UAS:ntr-mCherry)* zebrafish lines was used to label glomerular podocytes in larvae. Larvae were grown to three days post-fertilization and subjected to mild proteolysis with 1 mg/ml collagenase to disaggregate larval tissues. mCherry+ glomeruli were collected by pipetting under an epifluorescent stereoscope (Leitz M165) and washed 4x in HBSS with Calcium. Glomerular RNA was isolated from three glomerular isolation biological replicates with an RNAeasy kit (Qiagen), and poly(A) selected mRNA was used to prepare cDNA libraries (TruSeq Stranded mRNA Library Prep Kit), which were then subjected to illumina Hi-seq sequencing (50 bp paired end reads). Larval cDNA libraries of the same stage were prepared similarly and sequenced as a comparator for glomerular enriched gene expression.

### RNA-seq data processing

Paired-end FASTQ reads were processed with the nf-core/rnaseq pipeline (v3.26.0, Nextflow) using default parameters, against the *Danio rerio* GRCz11 genome and Ensembl release-115 annotation (Danio_rerio.GRCz11.dna_sm.primary_assembly.fa.gz, Danio_rerio.GRCz11.115.gtf.gz). The pipeline’s default workflow performed read trimming, STAR alignment, and Salmon quantification, yielding a merged gene-level count matrix (gene IDs, gene names, and raw counts per sample) used for all downstream analysis. Sequence data is deposited in GEO under accession GSE347631.

### Differential expression analysis (DESeq2)

Gene-level counts for the six embryo and glomerulus libraries were analyzed in DESeq2 (v1.48.2, R 4.5.1; design = ∼fraction, reference level = embryo). Genes were pre-filtered to those with at least 100 counts in two or more samples. Differential expression (glomerulus vs. embryo) was tested with the DESeq2 Wald test; log2 fold changes were shrunk with apeglm (v1.30.0), while Benjamini-Hochberg-adjusted p-values were retained from the unshrunk Wald model. Genes with padj < 0.05 were considered significant. Variance-stabilizing (VST) and regularized-log (rlog) transformations (blind = FALSE) were generated for visualization (PCA, volcano, and hierarchically clustered heatmap of significant genes) and as companion expression tables.

### Gene set enrichment analysis (GSEA)

Preranked GSEA (clusterProfiler v4.16.0) was performed on the full set of expressed, tested genes (n = 15,184), ranked by DESeq2 log2 fold change. Gene sets were drawn from Gene Ontology Biological Process terms (*D. rerio* annotations via biomaRt v2.64.0), filtered to a minimum/maximum gene-set size (GO: 10-5000 genes) and evaluated with Benjamini-Hochberg-adjusted p-values. Full bidirectional GO:BP results (both glomerulus- and embryo-enriched terms, all p-values) are reported in the output tables; corresponding summary figures are restricted to terms enriched in glomeruli (NES > 0, padj < 0.05) for clarity.

### Whole-mount in situ hybridization and Histology

Whole-mount *in situ* hybridization was performed as previously described (Thisse et al., 2004) with modifications (Kamei et al., 2015). Antisense RNA probes were generated from 500–800 bp cDNA blocks (Integrated DNA Technologies (IDT)) cloned into a pCRII-TOPO vector, linearized using the appropriate enzyme and transcribed using T7 RNA polymerase (New England Biolabs) in the presence of digoxigenin-labeled nucleotide triphosphates (Roche).

Embryos (72 hpf) were fixed in 4% PFA/PBS overnight at 4°C, washed in PBT (0.1% Tween®20/PBS), permeabilized with proteinase K (10 µg/mL in PBST; 20-45 min depending on the stage at room temperature) and underwent pre-hybridization washes in hybridization buffer (50% formamide, 5x SSC, 50 µg/mL heparin, 500 µg/mL tRNA, 0.1% Tween®20, pH 6.0) at 68°C. Embryos were hybridized with the digoxigenin-labeled probe. Probes were detected using anti-DIG-Alkaline Phosphatase (1:5000;Roche) and the NBT/BCIP chromogenic substrate (Boehringer Mannheim). After color development, embryos were fixed with 4% PFA, cleared with dimethylformamide, treated with hydrogen peroxide to bleach pigmented cells, and mounted in glycerol/PBS (1:1) to be imaged on a Leica MZ12 microscope equipped with a Spot Image digital camera. For histological analysis, stained embryos were dehydrated through a graded ethanol series (50%, 70%, 80%, 90%, 100%; 10 min each) and embedded in JB-4 plastic resin based on the manufacturer’s instructions (Electron Microscopy Sciences). Sections (7 µm) were obtained using a Leica RM2165 rotary microtome, mounted with Permount (Fisher Scientific), and imaged using a Nikon E800 microscope equipped with a Spot Insight CCD digital camera.

### CRISPR Guide RNA Design

A total of 25 guide RNAs (gRNAs) were designed to target 13 protein-coding genes (two guides per gene except for *ptprq*) (Supplemental Table S1). Guides were selected to target known functional domains of the protein or early exonic regions, preferentially outside of exon 1 to minimize nonsense-mediated decay or genetic compensation. Target sequences were obtained from the UCSC Genome Browser, and gRNAs were designed using the Alt-R™ CRISPR-Cas9 guide RNA design tool (Integrated DNA Technologies (IDT)). Guides with predicted on-target and off-target efficiency scores above 50 were prioritized. When possible, guides were selected in close proximity to allow PCR-based genotyping using flanking primers for PCR. gRNA and primer sequences for RTPCR used are listed in Supplemental table S1. Genomic regions were sequenced prior to injections to exclude gRNA sequence polymorphisms.

### Microinjections to Generate CRISPR G0 Mosaic Mutants and DNA Extraction

For injections, crRNA and tracrRNA (100 µM each) were annealed (95 °C for 5 min, then gradually cooled to 25 °C) to generate a 50 µM RNA duplex, stored at −80 °C until use. Cas9 protein (Cas9 Nuclease V3, IDT) was diluted to 25 μM in PBS. To prepare the Ribonucleoprotein (RNP) injection mix (10 μM), 2 μl of Cas9 protein (25 μM), 1 μl of gRNA duplex (50 μM), and 0.75 μl of nuclease-free water were incubated at 37 °C for 5 minutes. Injection buffer (final concentration: 0.1% phenol red, 100 mM KCl, 10 mM HEPES) was added prior to injection of the RNP mix. 4.6 nl of the 10 μM RNP solution was injected into one-cell stage embryos using a Nanoject II (Nanoliter 2000 Microinjector, World Precision Instruments) following standard methods (Hoshijima et al., 2019). Injected embryos were maintained at 28 °C in E3 medium. Target sites were amplified by PCR and sequenced to confirm CRISPR-induced mutations. Genomic DNA was extracted (Meeker et al., 2007) from individual 2 dpf larvae by alkaline lysis (50 mM NaOH, 95 °C for15 minutes) followed by neutralization with 1 M Tris (pH 8.0) and used for PCR amplification of target loci.

### Primer design and Fragment analysis

Primers were designed to generate 150–600 bp amplicons with the CRISPR target site centered within the fragment and intronic regions avoided when possible. An M13F adapter sequence (5′-TGTAAAACGACGGCCAGT-3′) was appended to the 5′ end of each gene-specific forward primer. A two-step PCR strategy was used to enable fluorescent labeling. In the first reaction, targeted regions were amplified using the M13-tagged forward primer and a gene-specific reverse primer. The resulting product was diluted (1:50) and used as a template for a second PCR with a 6-FAM-labeled M13 forward primer. For specific targets, nested PCR was employed to enhance specificity. In such cases, an additional initial amplification with outer primers was followed by a second round with nested primers. PCR reactions were carried out using Phusion High-Fidelity DNA Polymerase (Thermo Fisher Scientific). Cycling conditions were 98 °C for 3 min; 40 cycles of 98 °C for 30 s, 57 °C for 30 s, and 72 °C for 30 s; followed by 72 °C for 10 min.

Final fluorescent amplicons were analyzed by capillary electrophoresis (SeqStudio Flex, Thermo Fisher Scientific) for fragment size determination. Data analysis was performed using Peak Scanner software (Thermo Fisher) according to the CRISPR-STAT method(Carrington et al., 2015). Indel frequency was calculated as the sum of all mutant peak intensities divided by the total peak intensity (mutant + wild type).

### Sequencing and ICE Synthego

For Sanger sequencing, eight injected embryos per gene were used. Genomic DNA extraction and PCR amplification were performed as described above, using gene-specific primers without M13 labeling. PCR products were purified using the QIAquick PCR purification kit (Qiagen) and quantified with a Nanodrop before sequencing. Purified amplicons were subjected to Sanger sequencing. Editing efficiency and indel spectra were analyzed using the ICE (Inference of CRISPR Edits) tool (Synthego) by comparing chromatograms from injected samples to wild-type controls.

### Proteinuria assay, dextran injections

GFB integrity was assessed using a dual-dextran filtration assay based on a previously described method(Kotb et al., 2014). Larvae were anesthetized with 0.02% tricaine (Sigma-Aldrich) in E3 before microinjection. At 4 dpf larvae were injected retro-orbitally with a 1:1 mixture of 5mg/ml of 10 kDa rhodamine dextran and 500 kDa fluorescein (FITC) dextran. Approximately 23 nl of the solution was injected per larva using a Nanoject 2000 (World Precision Instruments). After injection, larvae were transferred to fresh E3 media to recover for 2 h, then screened under a fluorescence microscope (Leica M205FA) to confirm systemic distribution of the dextran. The successfully injected fish were kept at 28 °C for 48 h before imaging.

### Dextran injection imaging and quantification

The injected fish were euthanized with tricaine, fixed overnight at 4 °C in 4% PFA, washed in PBST (0.1% Tween20 in PBS), and mounted in 50% glycerol/PBS for imaging. The fish were mounted laterally on glass slides with a No.1.5 coverslip (Fisher brand). Images were acquired using a point scanning confocal unit (Zeiss LSM 980 confocal microscope, Carl Zeiss Microscopy) mounted in an Axio Examiner Z1 upright stand (Carl Zeiss Microscopy)and equipped with a Plan-Apochromat 10X/0.45 objective (Carl Zeiss Microscopy). Images were acquired using the Airyscan 2 detector in super resolution mode and controlled with ZEN Blue 3.1 software.

FITC-conjugated 500 kDa and tetramethylrhodamine-conjugated 10 kDa were excited by the 488 nm diode and the 561 nm DPSS lasers, respectively. Emission detection ranges were set to 499-557 nm for FITC and 573-627 nm for tetramethylrhodamine. Laser power and detector gain were adjusted based on histogram feedback to maintain signal intensities within the linear dynamic range, typically targeting 25–50%, to avoid saturation. Acquisition settings were kept constant within each experiment to allow quantitative comparison.

Z-stacks (10x objective) spanning proximal pronephric tubules were acquired using a Motorized Scanning Stage 130 × 85 PIEZO (Carl Zeiss Microscopy) mounted on Z-piezo stage insert WSB500 (Carl Zeiss Microscopy) at 16-bit depth and bidirectional scanning mode. Airy scan processing was performed using manual settings (processing strength = 1), and images were saved in CZI format for downstream analysis.

Fluorescence intensity of dextran reuptake was quantified in the proximal tubule using FIJI (ImageJ; version 1.47, National Institutes of Health). Regions of interest (ROIs) were manually outlined based on the 10 kDa dextran signal. For quantitative analysis of fluorescence intensities, quantification was performed using the formula for Corrected Total Cell Fluorescence (CTCF) (Ansari et al., 2013) CTCF = integrated density − (ROI area × mean background fluorescence). This formula gave us the CTCF of a single stack of a single channel.

The selected stack was the one that presented the largest area of the tubule. The background fluorescence was determined as the average of three independent regions per image. Glomerular filtration barrier integrity was expressed as the ratio of 500 kDa to 10 kDa dextran CTCF values. Data was normalized to controls within each experiment.

### Statistical analysis

Statistical analysis and graphs were performed using GraphPad Prism V 10.5.0. Values are presented as mean ± SD. Statistical significance was determined by ANOVA for multiple comparisons, while two groups were analyzed using Student’s 2-tailed unpaired t-test. Differences were considered significant with p-values < 0.05.

## Results

### The zebrafish glomerular transcriptome

We generated a bulk RNA seq transcriptome of micro-dissected zebrafish pronephric glomeruli at 2-3 dpf, the stage of pronephric kidney development when blood vessels sprouting from the dorsal aorta invade clusters of podocyte progenitors to generate the pronephric capillary tuft and glomerular filtration barrier (Drummond and Davidson, 2010). By 4 dpf, pronephric podocytes establish slit-diaphragms that contribute to the size-selective blood filtration barrier of the kidney (Kramer-Zucker et al., 2005). We chose bulk RNA sequencing of micro-dissected glomeruli over single cell approaches in order to maximize the sensitivity of transcript detection and better capture low-expressed genes and full transcripts.

Glomeruli from *Tg(podocin:gal4)* X *Tg(UAS:ntr-mCherry)* larvae were isolated by enzymatic digestion and purified by serial manual pipetting 3X under a fluorescent dissecting microscope (figure 1A,B). RNA extracted from 250 glomeruli and stage-matched whole larvae was subjected to bulk RNA-seq analysis. Differential expression analysis using DESeq identified 1,549 glomerular transcripts enriched more than 4-fold (>2 log2FC) over stage-matched whole larval transcriptomes and with an adjusted p-value of less than 0.01 (Supplemental figure 1, Supplemental Table S2).

**Figure 1.**
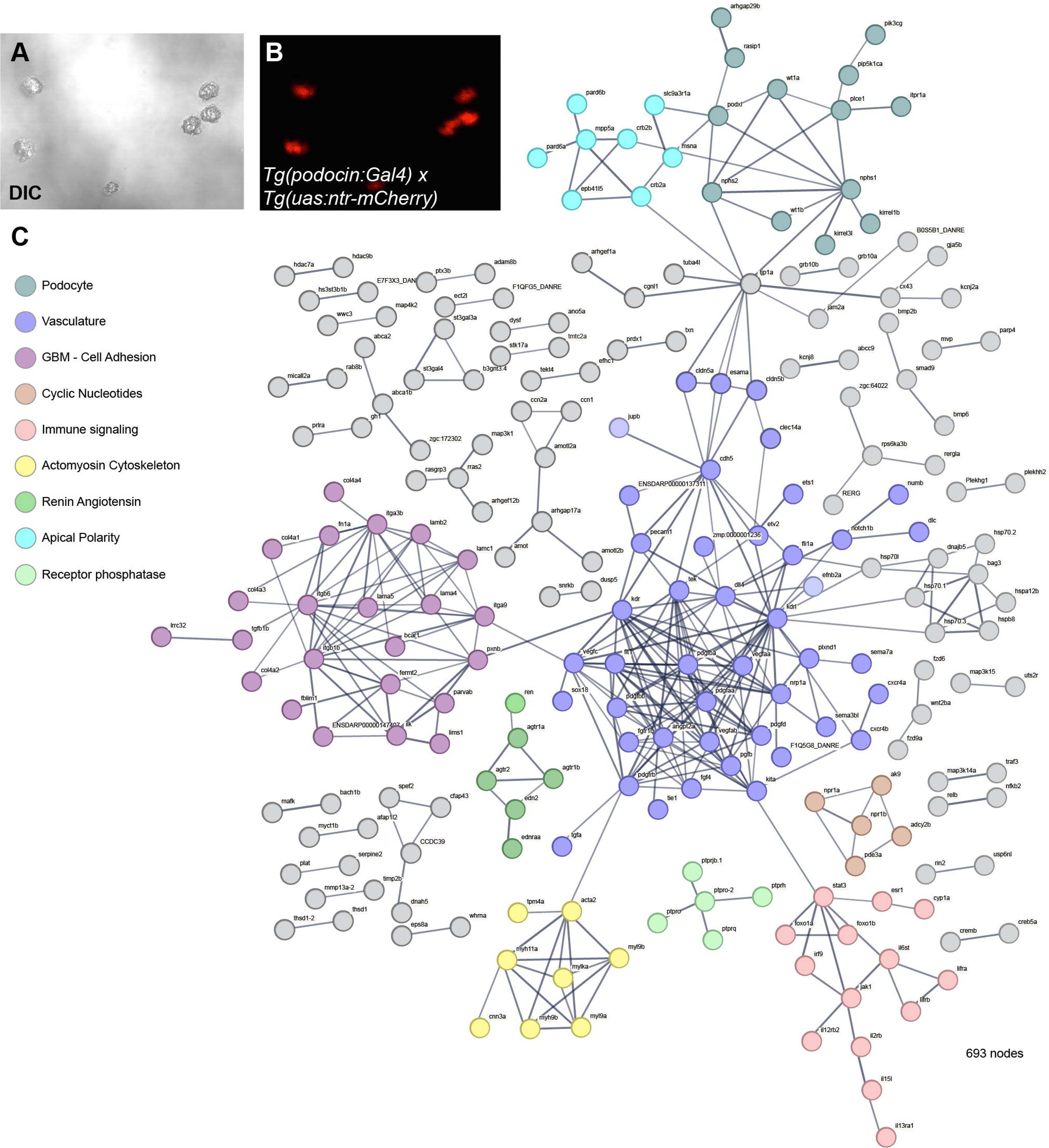
Glomerulus isolation and String interaction plot of proteins encoded by the zebrafish glomerular mRNAs. (A) DIC image of collagenase isolated 3 days post-fertilization (dpf) *Tg(podocin:gal4) X Tg(UAS:ntr-mCherry)* zebrafish glomeruli. (B) Fluorescence image of the same glomeruli as in A. (C) StringDB analysis plot of proteins encoded by glomerular mRNAs includes known podocyte and endothelial protein networks.

Ontology and gene set enrichment analysis revealed the expected enrichment of glomerulus development (GO:0032835) genes and blood vessel morphogenesis (GO:0048514) genes as well as GO terms relating to organ development (Supplemental figure 2; Supplemental Table S3). Protein-protein association mapping with the String Db (Szklarczyk et al., 2025) additionally identified gene networks regulating glomerular basement membrane formation, immune signaling, renin-angiotensin signaling, apical cell polarity, and actin cytoskeletal regulation (figure 1C). Of 498 genes enriched more than 4-fold in human podocyte single-cell RNA-seq data sets (KPMP, 2026), 124 (25%) were represented by orthologs or related genes (blastp e-value < 1e-120) in the zebrafish glomerular transcriptome (Supplemental Table S4).

### A CRISPR screen of glomerulus-enriched genes

To gain insights into zebrafish glomerular morphogenesis we conducted a limited G0 CRISPR screen, targeting genes highly enriched in developing zebrafish glomeruli that had not been characterized in human kidney development, rejecting candidates that were enriched in a previously established transcriptome of zebrafish endothelial cells ((Cannon et al., 2013), figure 1C, Supplemental table S2, S3). Of the remaining genes, we focused on genes either known to be expressed but uncharacterized in mammalian podocytes (*ptprq*) or whose function was related to cellular pathways known to play a role in glomerular morphogenesis (actin organization, cell-cell junctions, G-protein regulators). Thirteen genes (12 candidates plus *nphs*1 as a positive control) were selected for CRISPR targeting (Table 1). Glomerular enrichment was confirmed for *ptprq* and *fam49ba* by whole-mount in situ hybridization (Figure 2A,B) and podocyte enrichment of *ptprq* was demonstrated by histological sectioning of whole-mount in situ (Figure 2C).

**Figure 2.**
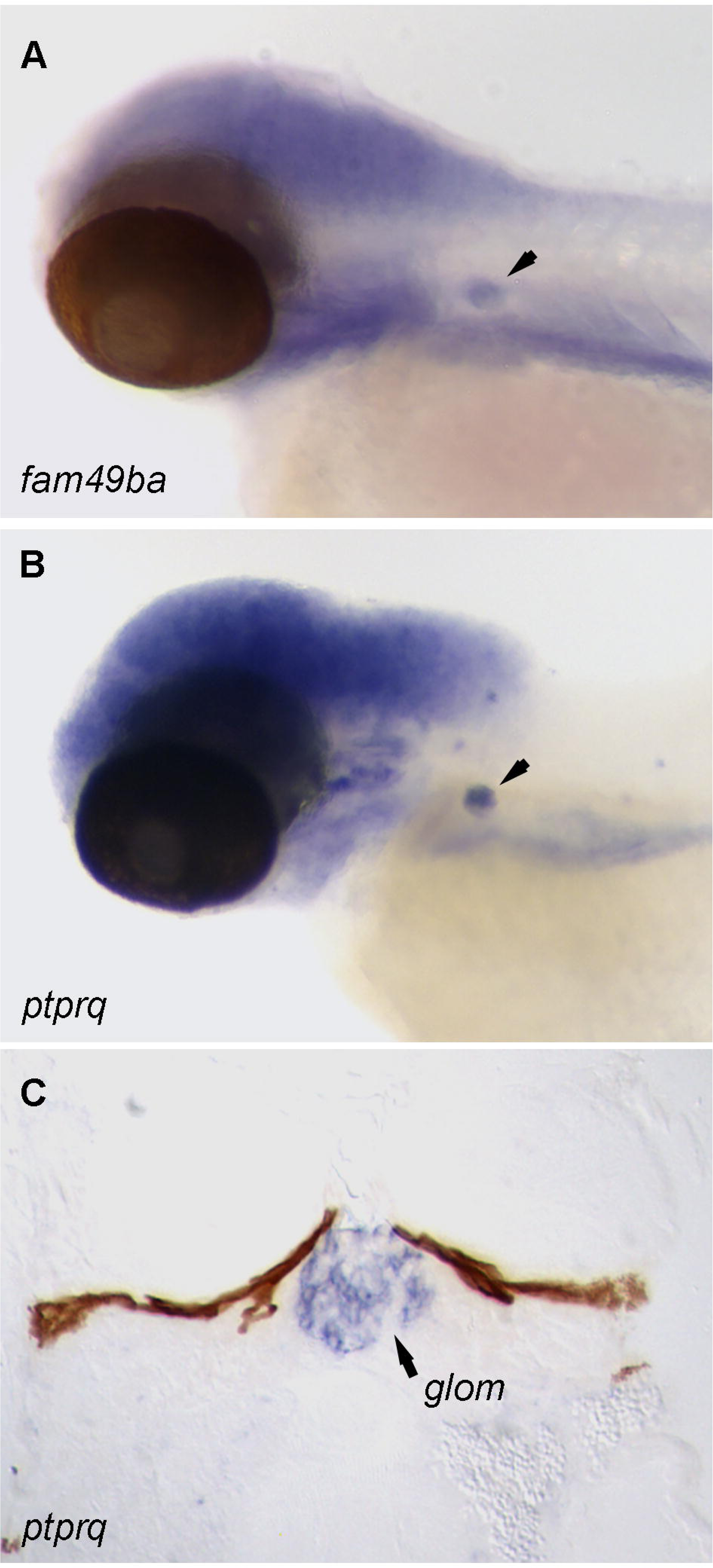
Podocyte-specific expression of glomerulus enriched mRNAs *ptprq* and *fam49ba*. Whole-mount in situ expression of *ptprq* (A) and *fam49ba* (B) in 3dpf zebrafish larvae. Arrowheads indicate pronephric glomerulus expression. (C) Histological section of *ptprq* in situ hybridization showing expression in 3 dpf glomerular podocytes.

**Table 1.** Glomerulus enriched genes selected for functional screening.

| **Gene ID** | **Symbol** | **Fold Enrichment** | **Abundance (baseMean)** |
|---|---|---|---|
| ENSDARG00000075720 | *il2rb* | 928.8 | 3694.3 |
| ENSDARG00000008849 | *ptprq* | 654.7 | 4007.0 |
| ENSDARG00000095873 | *nfxl1* | 635.9 | 418.6 |
| ENSDARG00000096757 | *frmd6* | 573.9 | 3639.4 |
| ENSDARG00000110020 | *lmo4* | 267.6 | 6011.6 |
| ENSDARG00000087832 | *bcl3* | 220.9 | 1116.3 |
| ENSDARG00000025866 | *ankdd1b* | 169.6 | 603.4 |
| ENSDARG00000095603 | *grid2ipb* | 61.7 | 3249.1 |
| ENSDARG00000016188 | *sbno2* | 51.7 | 10423.9 |
| ENSDARG00000039863 | *lifrb* | 39.1 | 3190.5 |
| ENSDARG00000077864 | *rasgrp3* | 33.3 | 1455.9 |
| ENSDARG00000020929 | *fam49ba* | 20.4 | 4632.7 |

Candidate glomerular genes were targeted using CRISPR-Cas9 RNP injections at the one-cell stage (Table 1, Figure 3A; see Methods). To assess CRISPR efficiency, genomic DNA from 48 hpf G0 Cas9 RNP injected larvae was analyzed by fragment analysis and Sanger sequencing to quantify indel formation. For each gene, PCR fragment analysis of uninjected control embryos showed a single peak product corresponding to amplification of the wild-type allele, while CRISPR-injected embryos displayed multiple PCR fragment peaks consistent with indel formation and successful mutagenesis (Supplemental Figure 3A,B). Sanger sequencing confirmed heterogeneous indels at the targeted loci in injected embryos, with no detectable alterations in uninjected controls (Figure 3B). Representative results from CRISPR-Cas9 targeting the *ptprq* gene show loss of the wild-type allele PCR fragment peak and generation of multiple indels at the targeted site (Supplemental Figure 2A). Quantification of editing efficiency demonstrated that all loci except *lmo4a* showed at least 75% loss of wildtype alleles, with several approaching 100% biallelic mutagenesis (Figure 3C, Supplemental Figure 3B). Hereafter, mosaic G0 CRISPR-Cas9 mutagenized larvae are referred to as “Crispants”.

**Figure 3.**
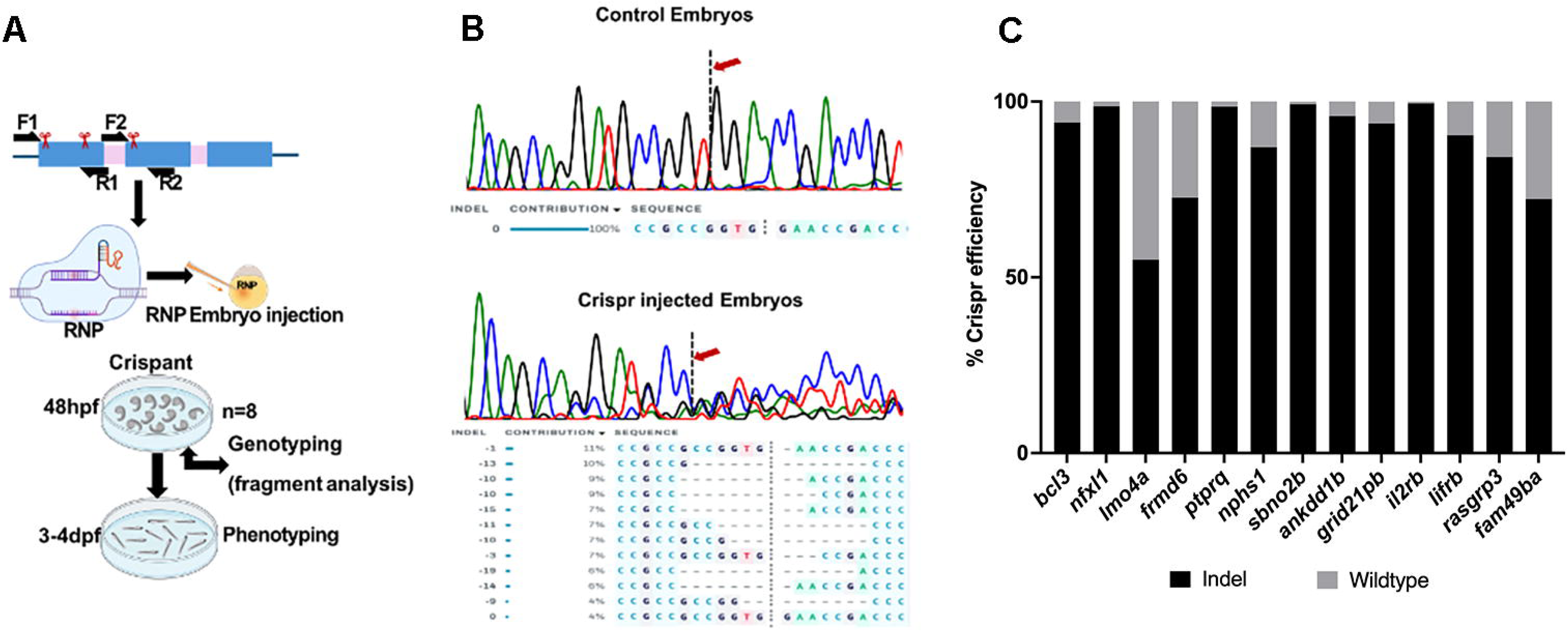
A CRISPR-Cas9 screen targeting zebrafish glomerular candidate genes. (A) Overview of the CRISPR-Cas9-based workflow used to target glomerular candidate genes in zebrafish. Synthetic gRNAs and Cas9 RNPs were injected at the one-cell stage. Embryos were genotyped at 48 hpf by PCR fragment analysis, followed by phenotypic screening at 4-6 dpf. (B) Representative Sanger sequencing traces confirming *ptprq* indel formation in CRISPR-injected embryos (bottom) compared to controls (top). (C) Quantification of CRISPR editing efficiency across 12 candidate genes and a positive control (*nphs1;nephrin*). Bar graph shows the proportion of embryos with detectable indels (black) versus no indel (gray), with most targets achieving mutagenesis rates over 70%.

To determine whether candidate genes were required for glomerular development and filtration barrier formation, we subjected crispant larvae to an in vivo dextran filtration assay. Larvae were injected with 10 kDa rhodamine dextran, which passes the glomerular filter and is taken up by endocytosis in the proximal tubules (Figure 4A; (Kotb et al., 2014; Kramer-Zucker et al., 2005)), along with 500 kDa FITC dextran which in healthy larvae is excluded from the kidney tubules by selective glomerular filtration. Glomerular barrier function was quantified as the ratio of 500 kDa to 10 kDa fluorescence intensity as a result of endocytosed tracer within the pronephric tubules (figure 4A). Control Cas9 injected larvae showed little or no passage of 500 kDa FITC dextran relative to 10 kDa rhodamine (Figure 4B) and showed a low fluorescent uptake ratio measurement (figure 4C). As expected, *nphs1/nephrin* crispants displayed a significant uptake of 500 kDa FITC dextran in proximal tubules and an increase in the 500 kDa:10 kDa ratio compared to controls (p < 0.001), indicating impaired filtration barrier selectivity (Figure 4B,C). Similar to *nphs1* targeted larvae, targeting *ptprq* and *fam49ba* also induced significant glomerular leakage of 500 kDa FITC dextran with uptake in proximal tubules (figure 4B,C), demonstrating that these genes have an essential function in glomerular morphogenesis. Among all genes screened for function this way, *nfxl1*, *lifrb, sbno2b, rasgrp3, bcl3*, *frmd6, fam49ba,* and *ptprq* crispants showed significant increases in 500 kDa:10 kDa dextran ratios compared to Cas9 controls (p < 0.05– 0.001). Fluorescence ratios for *lmo4a, il2rb, grid2ipb,* and *ankdd1b* crispants did not present significant differences from Cas9 controls, despite near complete mutagenesis (Figure 3C, Supplemental Figure 3B). These findings identify *nfxl1, lifrb, sbno2b, rasgrp3, bcl3, frmd6, fam49ba,* and *ptprq* as candidate genes for proteinuria and show they have an essential function in forming or maintaining the integrity of the zebrafish pronephric glomerular filtration barrier.

**Figure 4.**
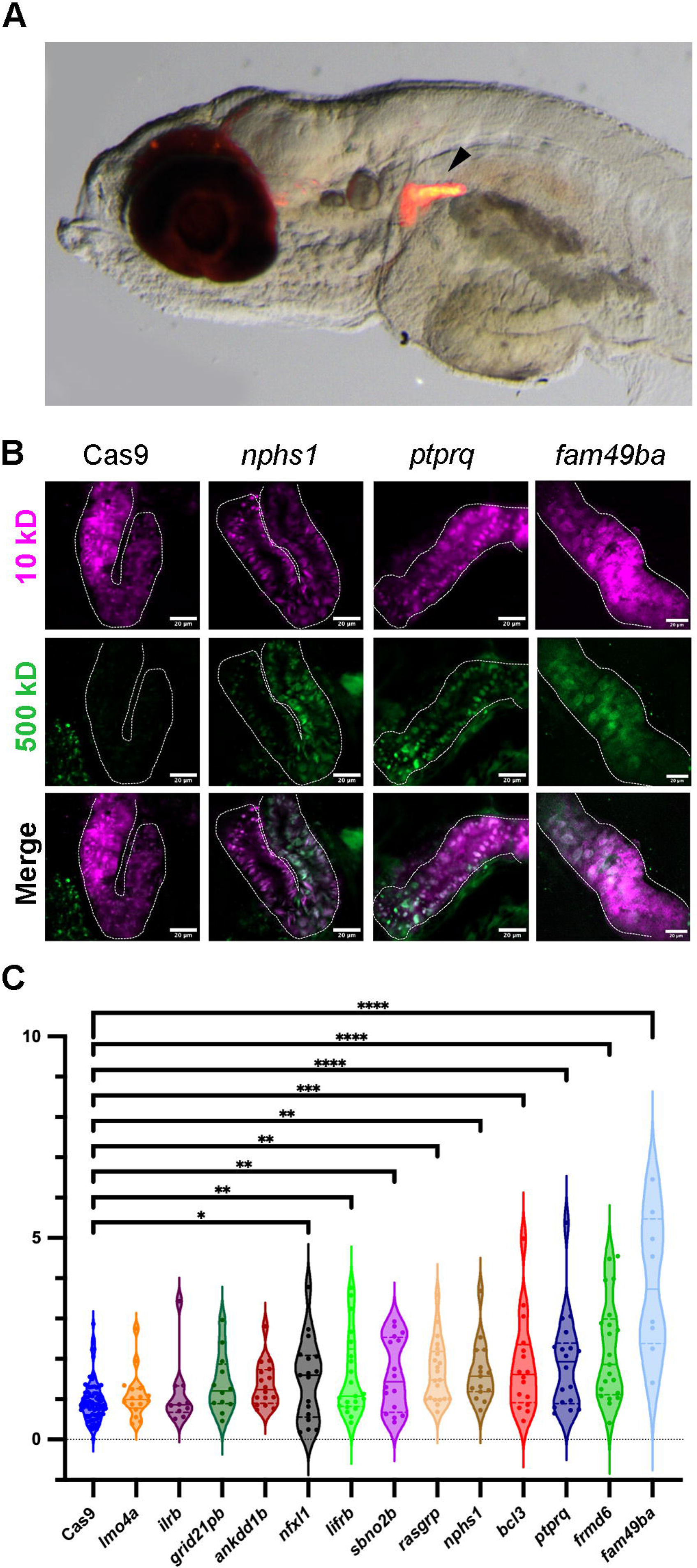
Detection of glomerular filtration defects by fluorescent dextran vascular injection. (A) Retro-orbital vascular injection of 10 kDa rhodamine dextran reliably labels endocytic vesicles of the pronephric proximal tubule (arrowhead). (B) Confocal imaging of proximal tubules in larvae injected with 10kDa rhodamine-dextran and 500kDa FITC-dextran. Cas9 injected embryos show 10kDa dextran uptake in proximal tubules but not 500kDa dextran. CRISPR mutagenesis of *nphs1*, *ptprq*, or *fam49ba* (*cyrib*) all show passage of 500 kDa dextran into proximal tubules, indicating a defective glomerular filtration barrier. Dotted line represents the area of fluorescence measurement. (D) Ratio quantification of 500kDa vs 10kDa dextran fluorescence signals for multiple CRISPR-Cas9 injected embryos identifies gene mutations causing statistically significant glomerular filtration defects.

## Discussion

Here we report transcriptomic profiling of developing zebrafish glomeruli coupled with in vivo CRISPR mutagenesis screen to identify novel regulators of kidney glomerular development and function. Comparative analyses revealed an overlap between the glomerular transcriptome of zebrafish and human podocytes, underscoring the evolutionary conservation of developmental programs governing nephrogenesis. Similar concordance of zebrafish and human podocyte expression profiles has been seen in single cell RNA seq study of adult zebrafish podocytes (Brown et al., 2021). It is likely that alignment of zebrafish genes to human orthologs in our work underestimates the similarity in gene expression because of differences in developmental stage (zebrafish larval vs. human adult), tissue complexity (whole glomeruli vs isolated podocytes), and experimental approach (bulk RNA seq vs. single cell RNA seq). Nonetheless, 124 orthologous genes that we identified highlight both known and yet to be characterized pathways in glomerular development.

We prioritized twelve genes enriched in zebrafish developing glomeruli that had not previously been associated with proteinuria, podocyte damage, or glomerulopathies. Functional perturbation using CRISPR-Cas9-mediated mutagenesis revealed that the disruption of eight genes, *fam49ba*, *frmd6*, *ptprq*, *bcl3*, *rasgrp3*, *sbno2b*, *lifrb*, and *nfx1l* resulted in significant proteinuria, as assessed by the dextran assay with three genes, *fam49ba, ptprq, and frmd6* showing the highest 500 kDa:10 kDa dextran ratios, indicated barrier disruption. In contrast, disruption of *il2rb, ankdd1b, grid2ipb,* and *lmo4a* did not induce detectable proteinuria.

Among the most significant mutational causes of proteinuria in our screen, *fam49ba* encodes Cyrib, a negative regulator of Rac1 (Fort et al., 2018; Ma et al., 2023), a small GTPase (Babelova et al., 2013; Blaine and Dylewski, 2020; Ma et al., 2023) critically involved in podocyte actin cytoskeleton organization and slit diaphragm stability (Asano-Matsuda et al., 2021). Disruption of *fam49ba* likely allows for uncontrolled Rac1 activity, leading to perturbation of the cytoskeletal architecture required for maintaining podocyte structure and filtration competence (Robins et al., 2017). *ptprq*, another highly significant screen hit encodes a podocyte-enriched putative cell adhesion protein linked to a phosphoinositide phosphatase domain, regulating pip2/pip3 levels. Phosphoinositide levels are known influence calcium signaling and actin dynamics, processes that are critical for foot-process architecture and podocyte morphogenesis (Djenoune et al., 2021; Gass et al., 2022; Goodyear et al., 2003; Jeon and Zinn, 2015; Pulido et al., 2013; Senju and Lappalainen, 2019). *frmd6*, a third high significance hit, encodes the FERM domain-containing protein Willin that functions upstream of Hippo signaling and regulates actin cytoskeleton dynamics. As Hippo signaling is required for normal organ development and regeneration, including the kidney, our findings support a potential role for *frmd6* in glomerular development (Angus et al., 2012; Chen et al., 2021; Gunn-Moore et al., 2005; Ishiuchi and Takeichi, 2011; Lara et al., 2025; Riley et al., 2022).

Remaining causes of glomerular leakage include mutation of *rasgrp3*, a Ras/Rap1 guanine nucleotide exchange factor expressed in the glomerulus. Rasgrp3 activates Rap1, which acts downstream of Nephrin to regulate podocyte integrin adhesion and slit diaphragm integrity. Mutation of *rasgrp3* may contribute to a proteinuria phenotype through a mechanism involving Rap1-integrin signaling (Maywald et al., 2022; Roberts et al., 2004; Stone, 2011; Yamashita et al., 2000). Several other genes in our screen function in regulation of inflammatory responses. *lifrb*, a receptor for leukemia inhibitory factor (Nicola and Babon, 2015; Xu et al., 2012), mediates cytokine signaling pathways involved in cell differentiation, survival and regeneration (Wang et al., 2023) and may be involved in cytokine signaling in kidney injury and repair (Liu, 2011). *bcl3* and *sbno2b* can both act as inhibitors of NFkB responses, limiting inflammation. Mutation in *bcl3* can result in unchecked NF-kB inflammatory signaling (Poveda et al., 2017). *Sbno2b* has been linked to STAT3, Notch-related signaling pathways, and NF-κB signaling (Brandstoetter et al., 2023; Coyle-Thompson and Banerjee, 1993; Majumdar et al., 1997), functioning as a feedback regulator of inflammatory responses (Brandstoetter et al., 2023; Grill et al., 2015; Syme et al., 2022). These results are consistent with a potential role for *bcl3* and *sbno2b* mutations in sensitizing glomerular cells to inflammatory responses initiated by injury or mutation (Jiang et al., 2023). Less is known about *nfxl1* which encodes a zinc finger transcription factor previously shown to be important in neural and epithelial development (Chintala and Katzenellenbogen, 2021). Although *nfxl1* has not been linked to kidney development, its identification in our screen suggests that it may play a role in glomerular gene regulation.

A subset of genes selected for our screen including *lmo4a*, a LIM domain-only transcriptional regulator, *il2rb*, primarily involved in immune signaling, *ankdd1b*, involved in protein binding and signal transduction, and *grid2ipb*, known for its role in regulating neuronal signaling, did not cause proteinuria, suggesting they are not essential for maintaining the glomerular filtration barrier under baseline conditions. Limitations of our study include the use of F0 crispants which introduces mosaicism and may influence phenotypic penetrance and variability. Although fragment analysis confirmed efficient mutagenesis, validation in stable mutant lines will be necessary to fully characterize gene-specific effects. Furthermore, the dextran filtration assay does not resolve the precise cellular compartment of deficiencies within the GFB, highlighting the need for complementary analyses such as transmission electron microscopy or single-cell transcriptomics.

## Conclusion

Our findings highlight the value of targeted functional screens of relatively understudied genes and pathways in kidney development. We show that several molecular pathways, from cytoskeletal regulation and calcium signaling to transcriptional control and inflammation can affect the development of the glomerulus. Although limited, our study opens avenues for developmental studies as well as translational exploration, as several pathways, including Hippo, Rac1, NF-kB, and cytokine signaling, are implicated in human glomerular diseases (Jiang et al., 2023; Robins et al., 2017; Sun et al., 2022).

## Acknowledgements

This research was supported by the National Institutes of Health (NIDDK 5UC2DK126021; I.A.D.) an NSF REU program grant (NSF DBI 2243416), and NIGMS IDeA Awards under P20GM103423, P20GM104318, and P30GM154610. Image collection for this manuscript was performed with the assistance of the MDI Biological Laboratory Light Microscopy Facility (RRID:SCR_019166). Animal husbandry was supported by the MDIBL COBRE comparative animal models core (P30GM154610).

## Figure Legends

**Supplemental Figure 1.** (A) Zebrafish 3 dpf glomerulus (Glom) and whole larvae (Emb) bulk RNA sequencing samples showing reproducible transcript abundance in three biological replicates. (B) Volcano plot showing genes differentially expressed between glomeruli and whole larvae of the same stage. Notable upregulated glomerular genes are marked, including the 12 candidate genes targeted in this screen.

**Supplemental Figure 2.** Gene set enrichment analysis. Preranked GSEA (clusterProfiler v4.16.0) was performed on the full set of expressed, tested genes (n = 15,184), ranked by DESeq2 log2 fold change. Gene sets were drawn from Gene Ontology Biological Process terms (*D. rerio* annotations via biomaRt v2.64.0), filtered to a minimum/maximum gene-set size (GO: 10-5000 genes) and evaluated with Benjamini-Hochberg-adjusted p-values.

**Supplemental Figure 3.** (A) Fragment analysis of control and *ptprq* gRNA / Cas9-injected embryos showing a reduction of the *ptprq* wild-type (WT) allele PCR product peak (arrow) and production of indel mutations in gene edited embryos. (B) Fragment analysis quantification of WT PCR fragment abundance for control Cas9-injected and Cas9/gRNA-injected embryos targeting the 13 genes surveyed in this screen.

