## Supplementary figures and images for "Novel regulators of zebrafish glomerular development"

### Supplemental figure 1

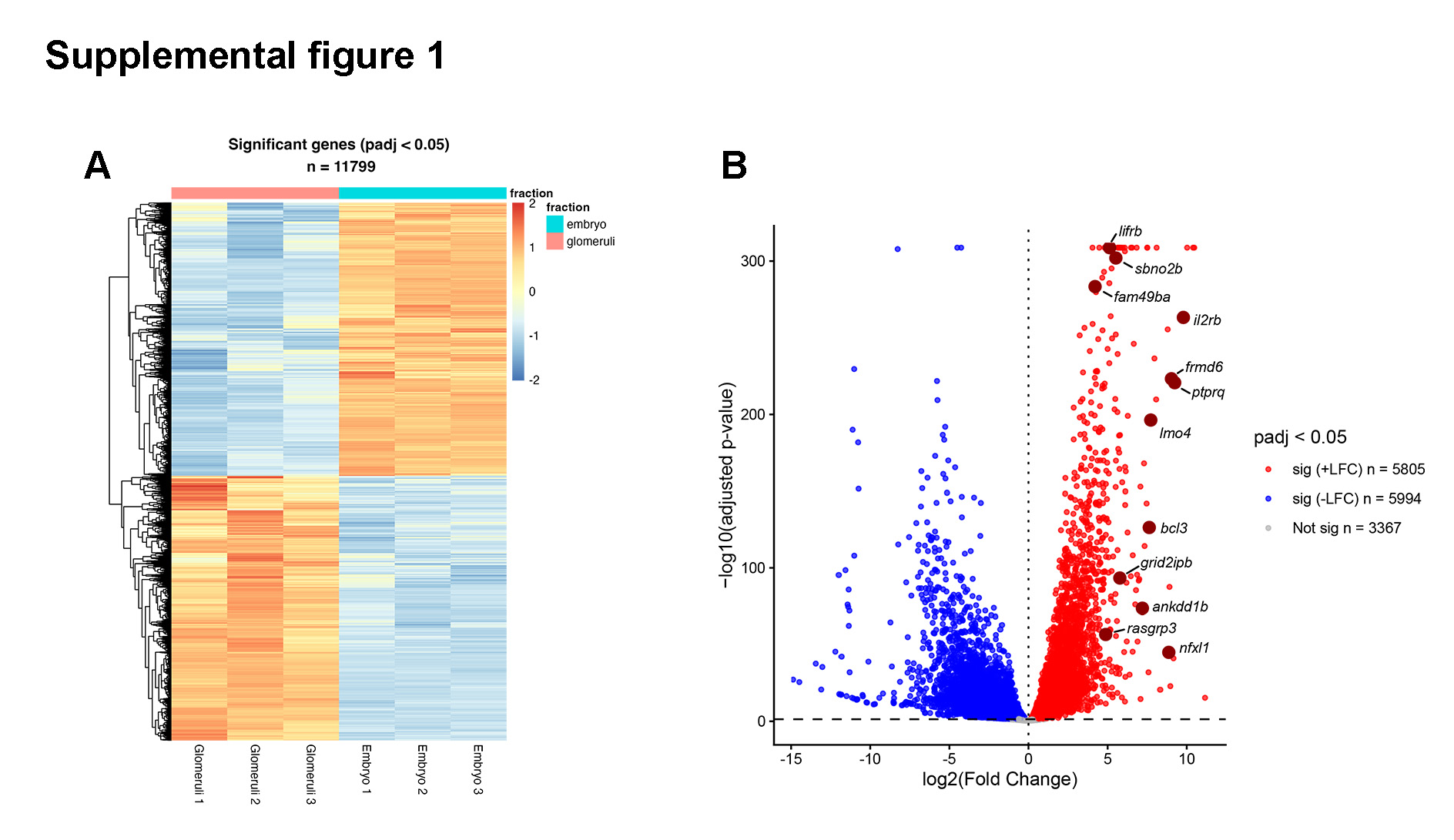

### Supplemental figure 2

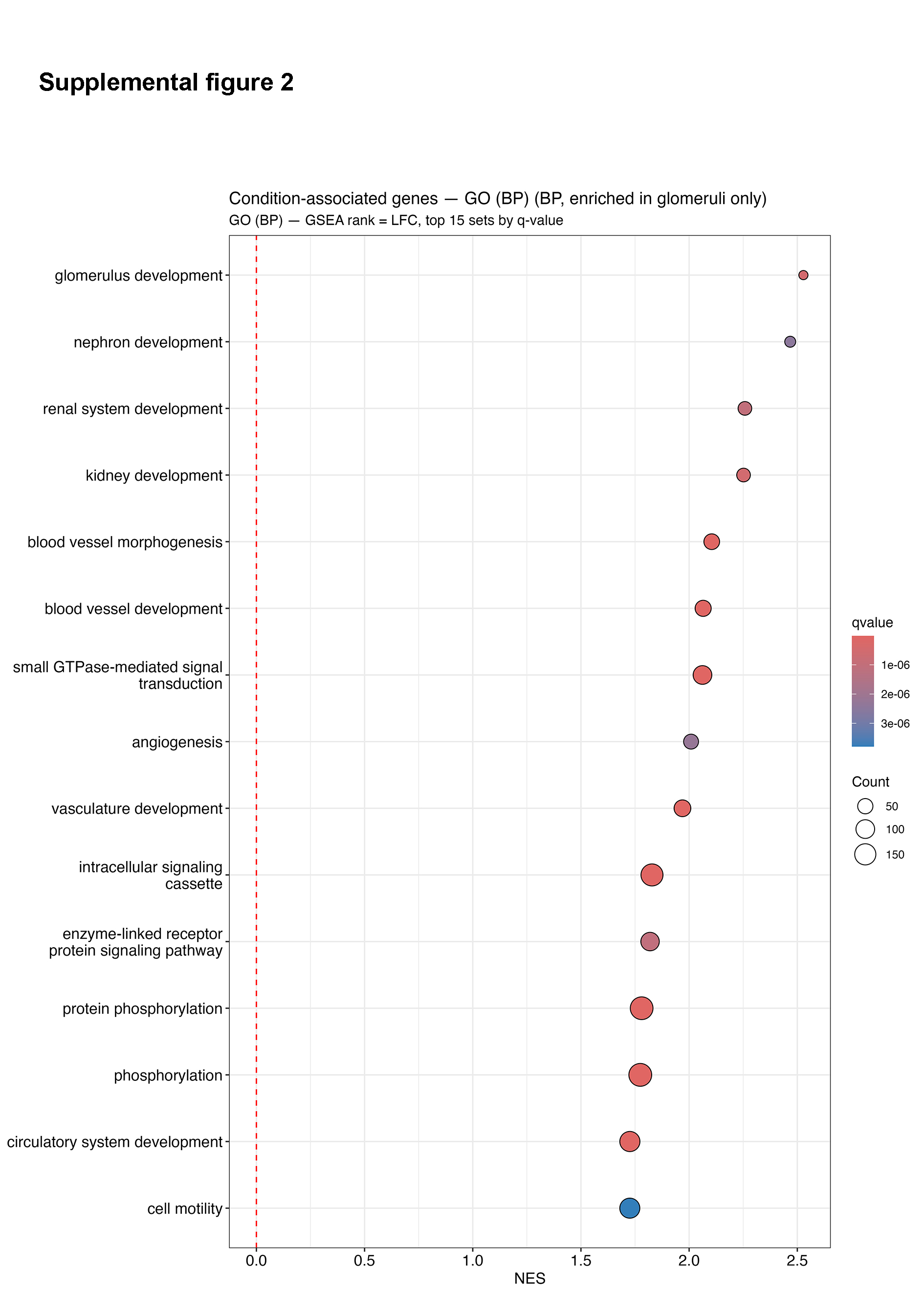

### Supplemental figure 3

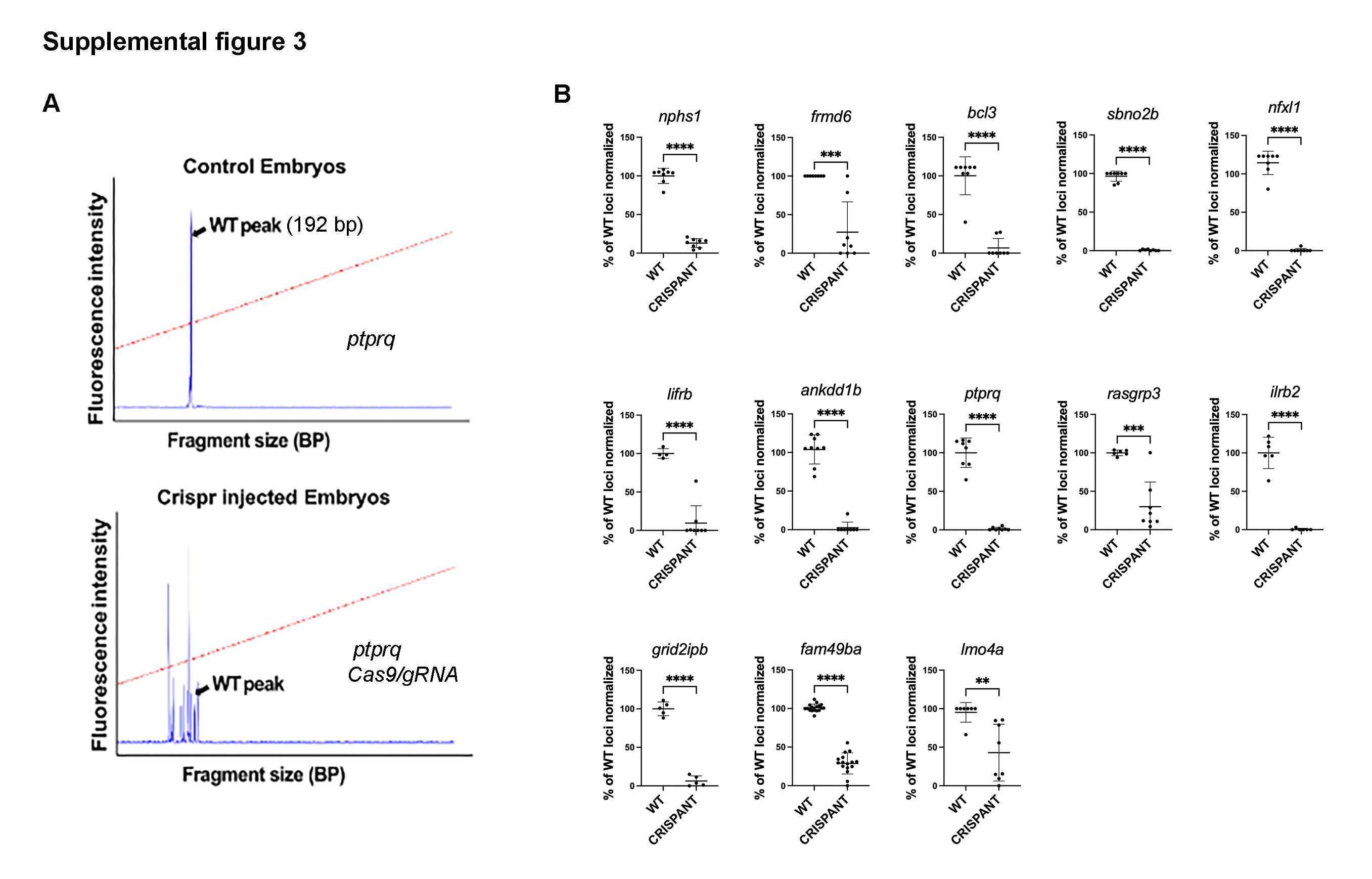
